# Uncovering a New Role of Dleu2/miR-15a/16-1 Cluster in Insulin Resistance and Obesity

**DOI:** 10.64898/2026.08.18.745519

**Authors:** Nitya Shree, Sunil Venkategowda, Mahua Choudhury

## Abstract

Obesity is a global epidemic characterized by metabolic dysfunction, with white adipose tissue playing a pivotal role in these processes. Noncoding RNAs, such as long non-coding RNAs (lncRNAs) and short non-coding RNAs (e.g., microRNAs), have been identified as an emerging class of regulatory molecules that can influence metabolic function. Here, the Dleu2/miR-15a/16-1 cluster (known as 13q14-Minimal Deleted Region, i.e., MDR), which encodes the lncRNA Dleu2 and miR-15a/16-1, a previously unrecognized player in metabolic function, is shown to contribute to obesity and insulin resistance. Using a combination of phenotypic and molecular approaches, this study establishes that MDR governs metabolic regulation for the first time. In a nutshell, this study identifies a new role of a lncRNA-miRNA cluster, previously implicated exclusively in cancer, in the regulation of obesity, thereby extending its biological significance beyond oncology.

**Graphical Abstract:** 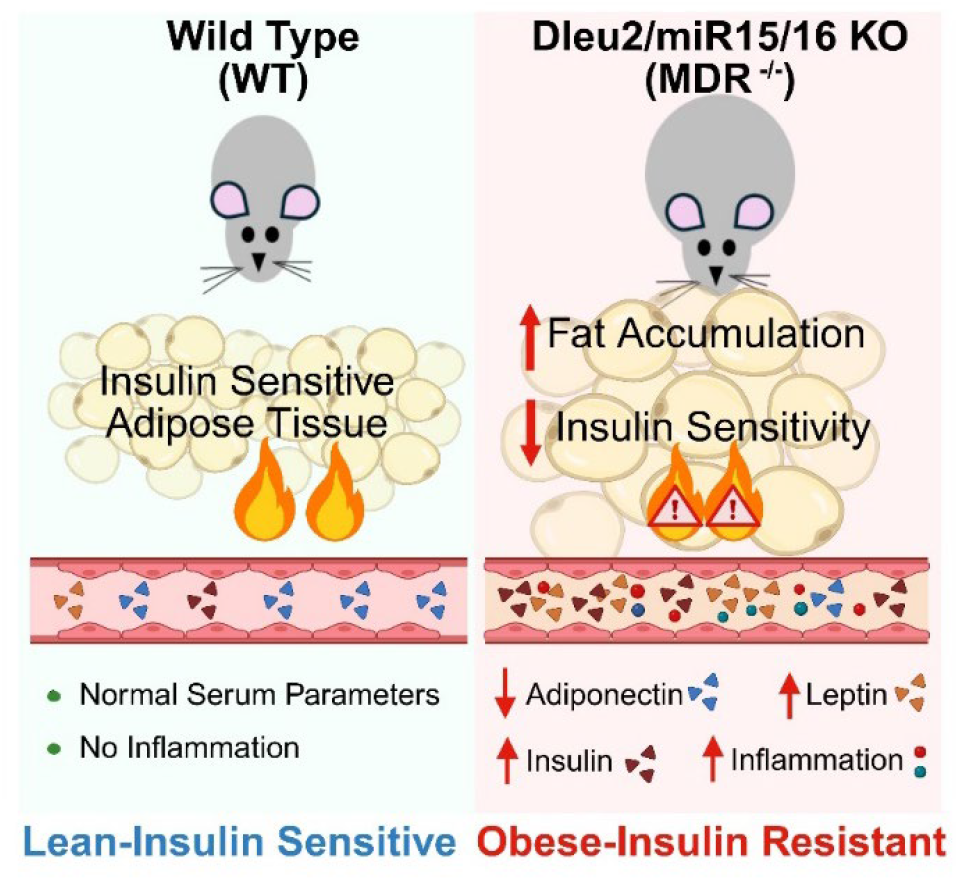

**Highlights:**

- Deletion of MDR contributes to obesity, insulin resistance, and impaired energy metabolism
- Loss of MDR reduces circulating adiponectin levels, indicating metabolic dysfunction
- MDR regulates satiety signaling in visceral adipose tissue and increases serum leptin levels
- MDR modulates several unrecognized new transcriptional regulators in obesity
- First evidence to establish the metabolic role of MDR beyond cancer biology

## Introduction

The prevalence of obesity has risen significantly worldwide, contributing to insulin resistance and type 2 diabetes (T2D) (Czech, 2017; Okunogbe et al., 2022). Adipose tissue is recognized as a dynamic metabolic organ, crucial for adipokine secretion and energy balance, that regulates metabolic homeostasis (Longo et al., 2019). Specifically, visceral adipose tissue (VAT) plays a pivotal role in this scenario (Oshakbayev et al., 2026). The role of VAT has long been underestimated as it has been considered merely a storage organ. However, it can become dysfunctional, leading to systemic metabolic compromise in the form of insulin resistance (IR) and eventually T2D (Longo et al., 2019; Macdougall et al., 2018). This dual epidemic has put a spotlight on adipose function as it is now recognized as an endocrine organ essential for regulating systemic energy homeostasis (Choe et al., 2016). More grave is that obesity-induced IR is also linked to a wide cluster of metabolic abnormalities, such as dyslipidemia, fatty liver disease, hypertension, coronary heart disease, and stroke (Deprince et al., 2020; Mostafa et al., 2025) . However, the mechanisms driving the origin of obesity are complex and not fully understood. Therefore, by revealing the early mechanisms of action in adipose dysfunction, there can be the formulation of future therapeutic strategies for these metabolic diseases.

During the past decade, noncoding RNAs (ncRNAs) have been linked to metabolic dysfunction (Lovell and Anguera, 2022). However, the discovery of new ncRNAs in VAT dysfunction is in its infancy. This study fills a significant knowledge gap regarding a lncRNA-miRNA cluster, which was originally only known in cancer biology. In the era of personalized and precision medicine, increasing our knowledge of adipose biology and identifying new early regulators will enable us to overcome the limitations of the traditional anthropometric indices of obesity. Notably, epigenetic changes are reversible, making them appealing targets for therapeutic and corrective interventions that are an urgent and unmet clinical need (Arguelles et al., 2016; Mahmoud, 2022)

More than a decade ago, Dalla-Favera’s group discovered the role of the Dleu2/miR-15a/16-1 cluster in the development of chronic lymphocytic leukemia (CLL) (Klein et al., 2010). The 13q14 minimal deleted region (MDR), which encompasses the Dleu2/miR-15a/16-1 cluster, is one of the most thoroughly studied tumor-suppressor loci in chronic lymphocytic leukemia and related malignancies (Xu et al., 2021; Zhang et al., 2022). To recapitulate the 13q14 deletion, a transgenic mouse model carrying the deletion of the minimal deleted region (MDR) was created (Klein et al., 2010; Zhang et al., 2024). However, the role of the MDR in obesity has never been investigated. Despite extensive research in the cancer arena, the potential influence of MDR on systemic metabolism, adipose tissue function, or energy balance remains uncharacterized. This gap is significant, as MDR encodes both a long non-coding RNA (Dleu2) and a microRNA cluster (miR-15a/16-1), suggesting its role as a regulatory hub at the intersection of transcriptional and post-transcriptional mechanisms.

In 2017, genome-wide association studies (GWAS) indicated that genetic variants in the human DLEU2 locus are associated with body weight, waist-to-hip ratio (-/+ adjusted to BMI), and hypertriglyceridemia, illustrative of their dysregulation in obesity and associated metabolic dysfunctions (Ng et al., 2017). In 2022, we provided the first experimental evidence that diet modulates Dleu2 and its downstream network, impacting metabolic function in the offspring, linking Dleu2 to metabolic disturbances over one or more generations (Zhang et al., 2022). A muscle-specific miR-16 knockout mice fed a high-fat diet was shown to impact metabolic and contractile properties, including glucose tolerance, insulin sensitivity, muscle contractile function, protein anabolism, and mitochondrial network health in a sex specific manner (Lee et al., 2016; Lim et al., 2022). On the contrary, in a small clinical study, miR-15a-5p was identified as a predictive plasma biomarker of cardiometabolic disease (Ramzan et al., 2020). However, the involvement of the Dleu2/miR-15a/16-1 cluster in whole-body metabolic regulation remains undefined, leaving a critical mechanistic gap in the field. Now, we are the first group to extensively characterize this complex MDR for its role in obesity and insulin resistance. This study provides a comprehensive metabolic characterization of this cluster utilizing a whole-body knockout of miR-15a/16-1 (MiR^-/-^) and MDR (MDR^-/-^) from Dalla-Favera’s group (Klein et al., 2010).

## Results and Discussion

### MiR^-/-^ and MDR^-/-^ mice exhibit increased fat accumulation, contributing to obesity

MiR^-/-^ and MDR^-/-^ mice were fed a standard chow diet for 32 weeks. Phenotypic assessments were performed at multiple timepoints during the experimental period (Figure 1A). Body weight was measured weekly, revealing a significant increase in weight gain in the MDR^-/-^ mice when compared to both WT and MiR^-/-^ mice, beginning at 6-weeks and 8-weeks of age, respectively. In contrast, MiR^-/-^ mice showed a significant increase in body weight, beginning at 23-weeks when compared to the WT group (Figure 1B). EchoMRI analysis revealed that the increased weight in both MiR^-/-^ and MDR^-/-^ mice was primarily attributed to significantly higher fat mass, whereas lean mass remained comparable across all groups (Figure 1C). Magnetic resonance imaging (MRI) of white adipose tissue (WAT) showed fat tissue compartment segmented into visceral WAT (VAT) and subcutaneous WAT (SAT) (Figure 1D). The distribution of both VAT and SAT was visibly higher in MDR^-/-^ than in WT and MiR^-/-^. When euthanized at 32 weeks, both knockout groups were evidently obese relative to WT (Figure 1E). Interestingly, MDR^-/-^ mice exhibited excessive fat accumulation compared to MiR^-/-^ mice (Figure 1F, S1A). Consistently, MDR^-/-^ mice also showed significantly higher WAT weight than WT and MiR^-/-^ mice (Figure 1F, S1E). Since VAT is recognized as the most metabolically active WAT (Dadson et al., 2026; Tchernof et al., 2006) we investigated if the relative increase in VAT mass was accounted for by increased adipocyte numbers (hyperplasia) or increased adipocyte size (hypertrophy). We observed a significant decrease in adipocyte numbers per field in both MiR^-/-^ and MDR^-/-^ mice, compared to WT mice with visibly hypertrophied lipid droplets (Figure 1G). Quantification also revealed a 2-fold increase in the MiR^-/-^ mice lipid droplet size, whereas a 2.2-fold increase in lipid droplets was documented in MDR^-/-^ mice (Figure 1H). Collectively, these findings confirm that the absence of MDR contributes to severe obesity with a more pronounced effect than the loss of miR-15a/16-1 alone by increasing fat mass accumulation even under a standard chow diet.

**Figure 1.**
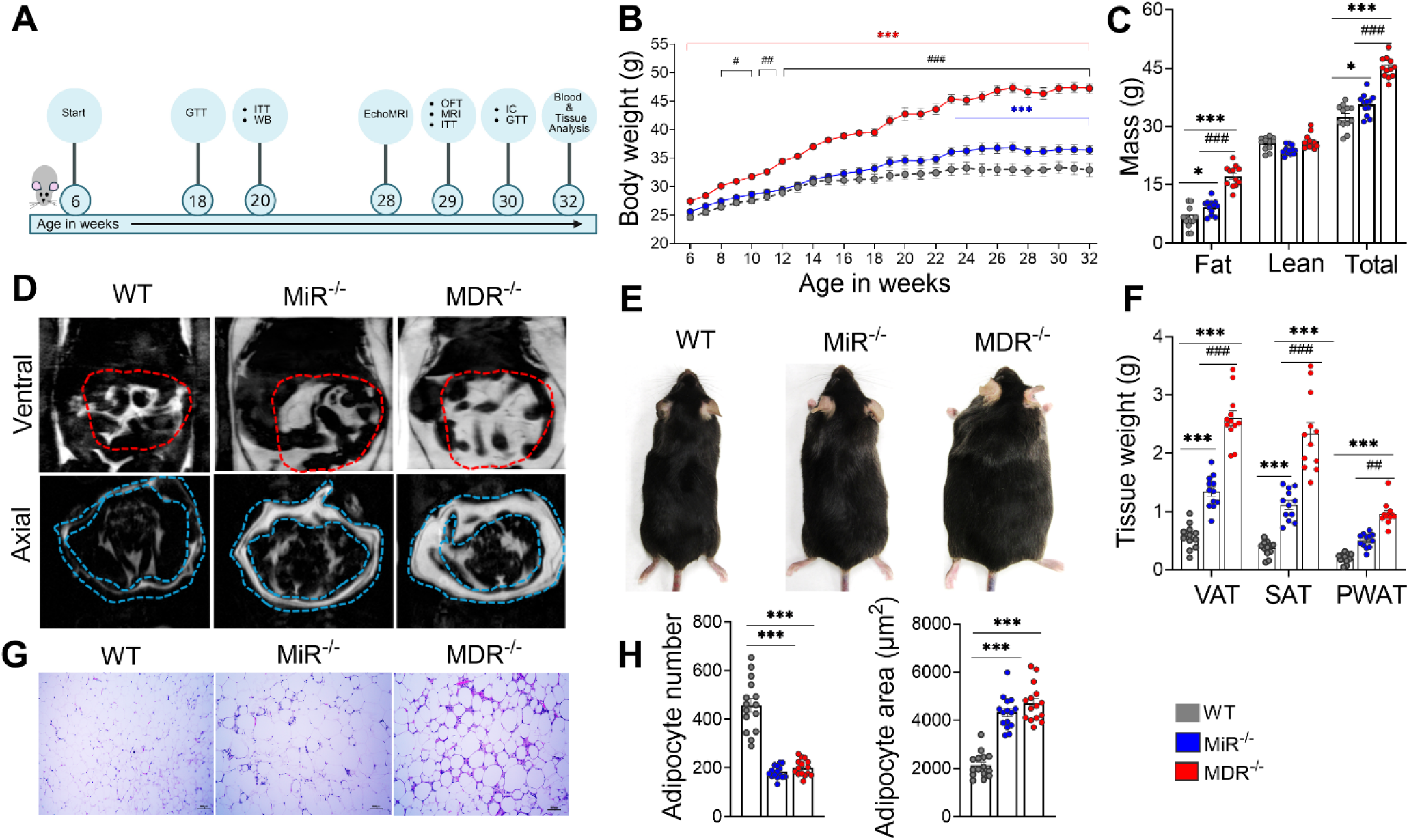
Whole-body MiR and MDR deletion contributes to obesity. (A) Schematic representation of study timeline (GTT: glucose tolerance test, ITT: insulin tolerance test, WB: western blotting, OFT: open field test, MRI: magnetic resonance imaging, IC: indirect calorimetry). (B) Body weight time course from 6 weeks to 32 weeks of age (*n*=12 per group). (C) EchoMRI at week 28 (*n*=12 per group). (D) Representative image of ventral and axial MRI of VAT (encircled red) and SAT (encircled blue) at week 29 (*n*=2 per group). (E) Representative image of the 32-week-old mice. (F) Fat depot (VAT, SAT, and perirenal white adipose tissue (PWAT) weight in mice at 32 weeks (*n* = 12 per group). (G) Representative H&E staining image of VAT sections (*n* = 3 per group). Scale bar, 100 μm. (H) Quantification of lipid droplet size VAT (*n*=3 per group) area. Data shown as mean ± SEM control. P values were calculated by Two-way ANOVA with Tukey’s post-test, except figure H, which was One-way ANOVA with Tukey’s post-test. \**p*<0.05,\*\**p*<0.01, and \*\*\**p*<0.001 represent significance compared to WT. MiR^-/-^ versus MDR^-/-^ is denoted as # *p*<0.05, ## *p*<0.01, and ### p< 0.001.

### MDR deletion drives insulin resistance and obesity-induced metabolic dysfunction

To determine the impact of MiR and MDR deletion on metabolic homeostasis, we performed glucose tolerance test (GTT) and insulin tolerance test (ITT) at week 18 and week 20, respectively. MDR^-/-^ mice showed a pronounced impairment of glucose tolerance compared to WT that could still be observed, albeit more mildly, in MiR^-/-^ mice (Figure 2A). In ITT, MiR^-/-^ and MDR^-/-^ mice displayed significantly impaired glucose suppression compared to WT mice (Figure 2B), indicating a profound insulin resistance phenotype. Both MiR^-/-^ and MDR^-/-^ mice retained their obese insulin-resistant phenotype until week 30 (Figure S1B, S1C). To directly assess the impairment in insulin signaling, we measured Akt2 phosphorylation (Ser473), which is the hallmark of obesity-induced insulin resistance in VAT. A significant downregulation in pAkt2/Total Akt was observed in MiR^-/-^ mice and MDR^-/-^ mice, confirming the dysregulation of insulin signaling (Figure 2C, S2A). Consistent with this, fasting serum insulin levels were also significantly higher in both groups when compared to WT (Figure 2D). Of note, MDR^-/-^ mice showed significantly higher insulin levels than the MiR^-/-^ mice. Both MDR^-/-^ and MiR^-/-^ mice had a higher HOMA-IR index than the WT mice, further supporting systemic insulin resistance (Figure S1D). Similar to insulin level, MDR^-/-^ mice showed significantly higher HOMA-IR index than MiR^-/-^ mice.

**Figure 2.**
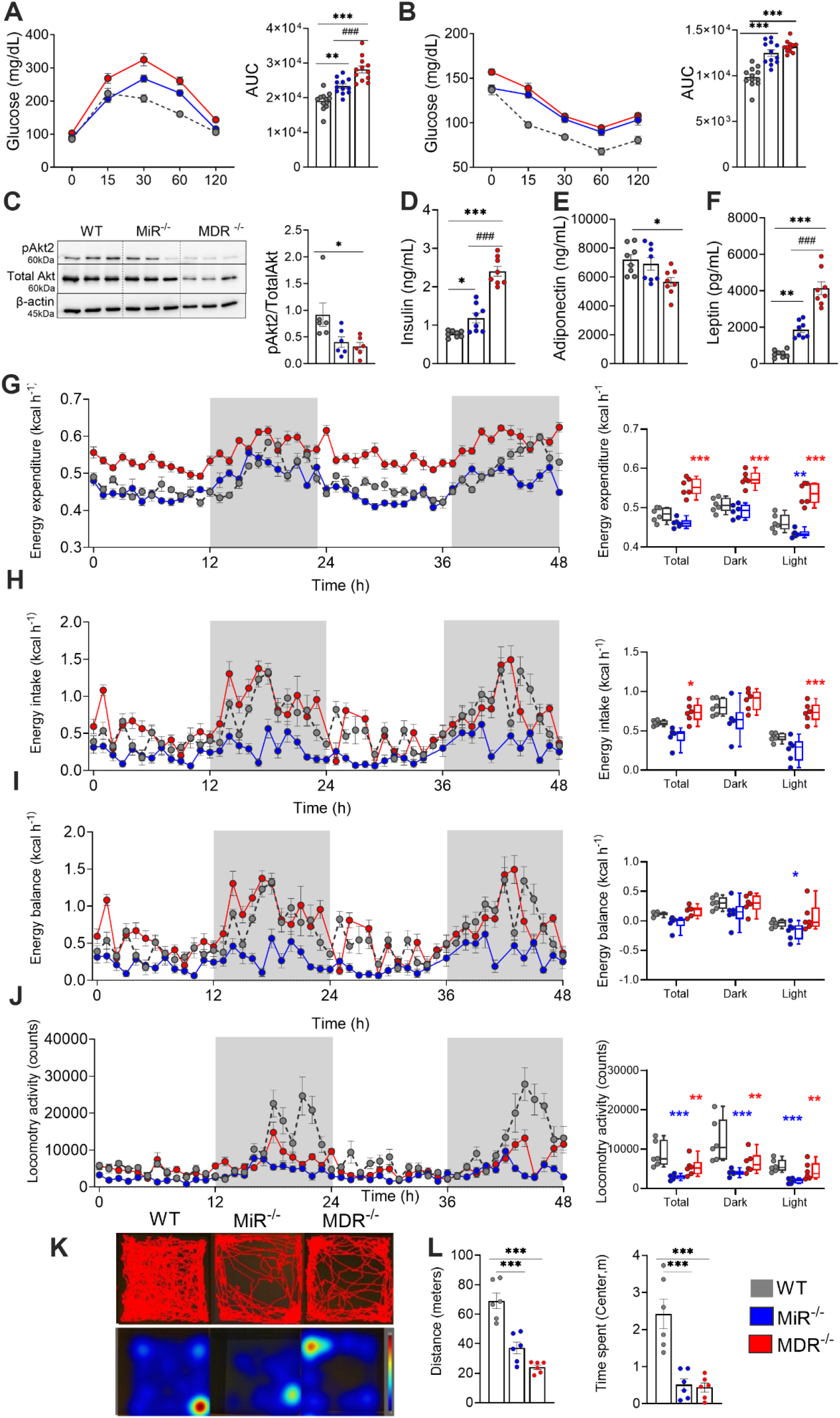
Whole-body MiR and MDR deletion are associated with insulin resistance and metabolic impairment. (A) Intraperitoneal glucose tolerance test (*n*=12 per group) at 18 weeks, and quantification of the area under the curve (AUC). (B) Insulin tolerance test (*n*=12 per group) at 20 weeks, and quantification of the area under the curve (AUC). (C) Representative western blot image of insulin signaling marker (*n*=3 per group) and quantification (*n*=6 per group). Serum biochemistry (*n*=8 per group), (D) Serum insulin, (E) Serum adiponectin, (F) Serum leptin. Indirect calorimetry for 48h (*n*=6 per group), (G) Energy expenditure. (H) Energy intake. (I) Energy balance. (J) Locomotory. Line graphs (G-J) show representative 48 h trace means and error bars ± SEM, and bar graphs show 48 h means and error bars ± SEM. (K) Representative track-plots obtained for each experimental group in Ethovision XT for the Noldus open field test. (L) Distance travelled and time spent in the center. Data shown as mean ± SEM control. P values were calculated by one-way ANOVA with Tukey’s post-test. \**p*<0.05,\*\**p*<0.01 and \*\*\**p*<0.001. MiR^-/-^ versus MDR^−/−^ is denoted as ### p< 0.001.

Previous work reported that the muscle-specific miR-16 knockout mice developed only a mild ITT slope, which was returned to the WT level after 60^th^ minute of ITT without any alteration in the pAkt (Ser473) levels (Lim et al., 2022). Our study revealed that the whole-body MiR^-/-^ mice exhibited impaired insulin tolerance until the 120^th^ minute of ITT and remained significantly higher than the WT level. Furthermore, in VAT, both MiR^-/-^ and MDR^-/-^ mice showed dysregulated insulin signaling pathway at the transcriptional (RNAseq gene expression, Figure 3) as well as translational level (Figure 2C), suggesting that VAT may be the primary regulator of systemic IR.

**Figure 3.**
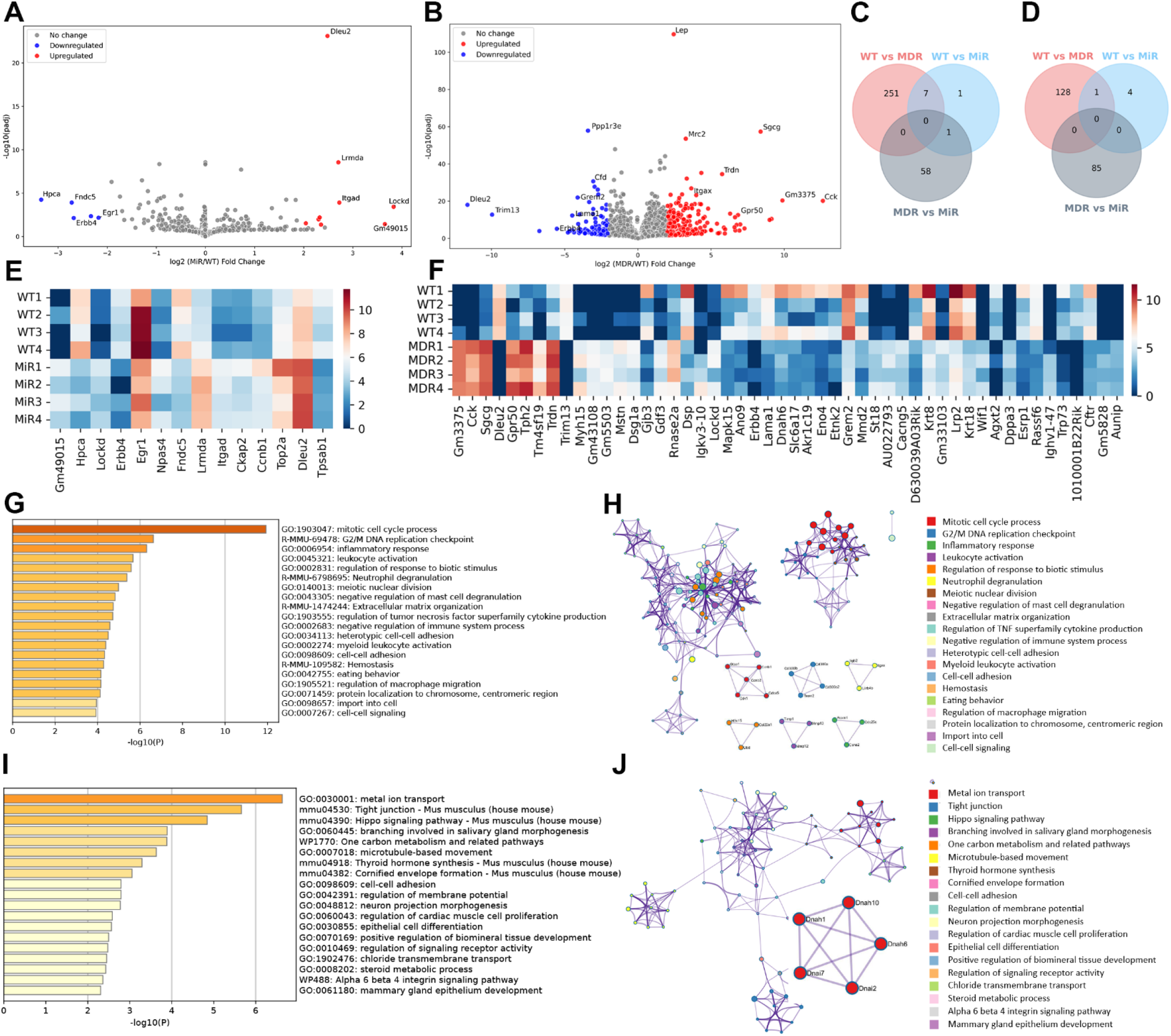
Transcriptomic and protein level alteration in VAT (n=4 per group). Volcano plot of differentially expressed genes. The y-axis indicates the -log_10_ (padj) and the x-axis indicates the log_2_ FC. Red circles represent upregulated genes (log_2_FC > 2, padj < 0.05) and blue circles indicate downregulated genes (log_2_FC < -2, padj < 0.05). Non-significant genes are shown as gray circles. (A) Volcano plot of WT vs MDR^-/-^. (B) Volcano plot of WT vs MiR. Venn Diagram showing uniquely expressed, (C) upregulated, and (D) downregulated genes. (E) Heatmap of top 25 upregulated and downregulated genes in WT vs MDR, (F) Heatmap of top upregulated and downregulated genes in WT vs MiR, (G) Metascape analysis of upregulated DEGs in MDR^-/-^ mice. (H) Protein-protein interaction enrichment analysis for upregulated genes. (I) Metascape analysis of downregulated DEGs in MDR^-/-^ mice. (J) Protein-protein interaction enrichment analysis for downregulated genes.

Serum profiling of obesity markers such as adipokines, lipids, and liver enzymes revealed a significant alteration in both MiR^-/-^ and MDR^-/-^ mice when compared to the WT mice, indicating broad metabolic disruption (Figure. 2 E-F, Table 1). Specifically, the MDR^-/-^ mice exhibited markedly reduced levels of adiponectin (p<0.05) and elevated levels of leptin (p<0.001) compared to WT mice, indicating disrupted adipokine secretion associated with obesity. In addition, both MiR^-/-^ mice and MDR^-/-^ mice had significantly altered alkaline phosphatase (ALP), high-density lipoprotein (HDL), low-density lipoprotein (LDL), non-esterified fatty acid (NEFA), and tumor necrosis factor-alpha (TNFα) compared to WT mice, reflective of systemic inflammation and metabolic dysregulation. There were no significant changes in other serum biochemistry parameters, including albumin, alanine aminotransferase (ALT), aspartate aminotransferase (AST), total cholesterol (TC), bilirubin, and triglyceride (Table 1). Many other parameters remained unaltered in MiR^-/-^ mice, as opposed to MDR^-/-^, when compared to WT. To emphasize, the impact has been more significant in MDR^-/-^ mice. Furthermore, complete blood count (CBC) analysis revealed a higher lymphocyte count in MDR^-/-^ mice compared to WT mice, suggesting chronic low-grade inflammation (Table 1). Overall, this again indicates that the effect of MDR^-/-^ is more profound than only MiR^-/-^ driving the IR or inflammation or obesity. To determine whether MiR or MDR deletion contributed to dysregulated whole-body energy metabolism, we performed indirect calorimetry followed by data analysis as per recently updated guidelines to preclinical indirect calorimetry experiments (Banks et al., 2025). Energy expenditure was reduced in MiR^-/-^ mice when compared to WT mice during the light period (Figure 2G). In contrast, MDR^-/-^ mice demonstrated significantly increased energy expenditure when compared to WT mice across both the dark and light periods. Energy intake showed similar patterns as the energy expenditure in both MiR^-/-^ and MDR^-/-^mice (Figure 2H), demonstrating that MDR^-/-^ mice have both increased energy intake and energy expenditure, contributing to positive energy balance and more weight gain as opposed to that observed in MiR^-/-^ mice (Figure 2I) (Banks et al., 2025). Furthermore, MiR^-/-^ mice showed a decreasing trend of O_2_ consumption as well as CO_2_ production only during the light cycle. In contrast, MDR^-/-^ mice showed a significant increase in both parameters when compared to WT in both dark and light cycles (Figure S2B, S2C). Overall, this suggests that the divergence in metabolic responses in MDR^-/-^ mice was stronger compared to that in MiR^-/-^ mice. All the groups maintained a consistent respiratory exchange ratio (RER), indicative of equal substrate utilization across day and night periods (Figure S2D). Both MiR^-/-^ and MDR^-/-^ mice showed reduced locomotor activity when compared to WT mice, confirming their obese phenotype (Figure 2J). In addition, the Noldus open field test demonstrated significantly reduced activity in MiR^-/-^ mice and MDR^-/-^ mice (Figure 2K, 2L). In a nutshell, the above outcome indicated that the MDR^-/-^ makes a greater contribution to insulin resistance, inflammation, and weight gain compared to the MiR^-/-^. Overall, this suggests that the divergence in metabolic responses in MDR^-/-^ mice was stronger compared to that in MiR^-/-^ mice. Noteworthy to mention that even with suppression of EE and no increase in EI, MiR^-/-^ mice consistently increased their body weight (Fig.1B). This again reminds us that a unifying concept of energy balance or weight gain cannot be used for personalized therapeutic strategy. Future studies in humans could investigate whether this could be associated with clinical advantages such as a reduction in tiredness, stress levels, and weight loss.

**Table 1.**
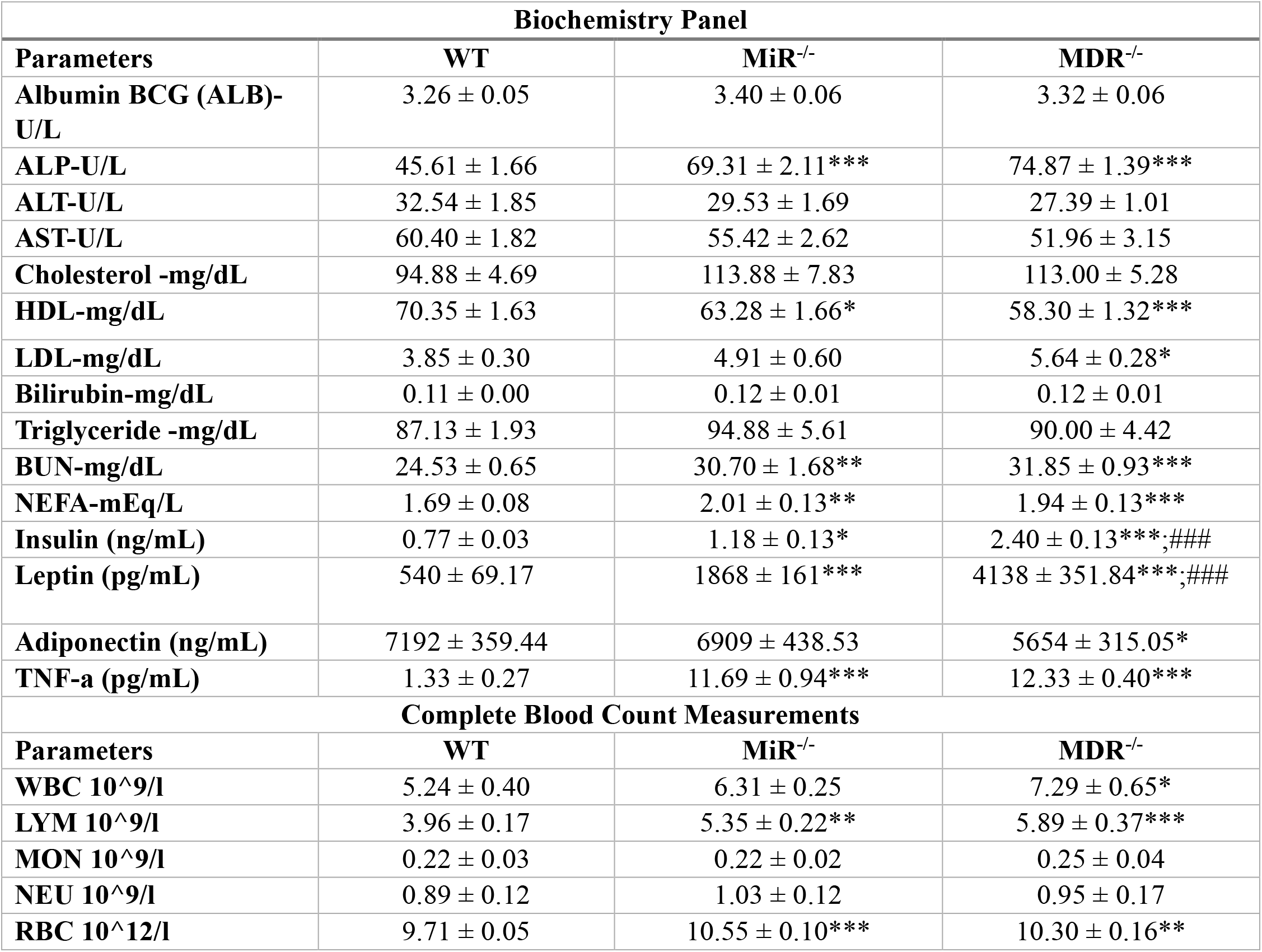
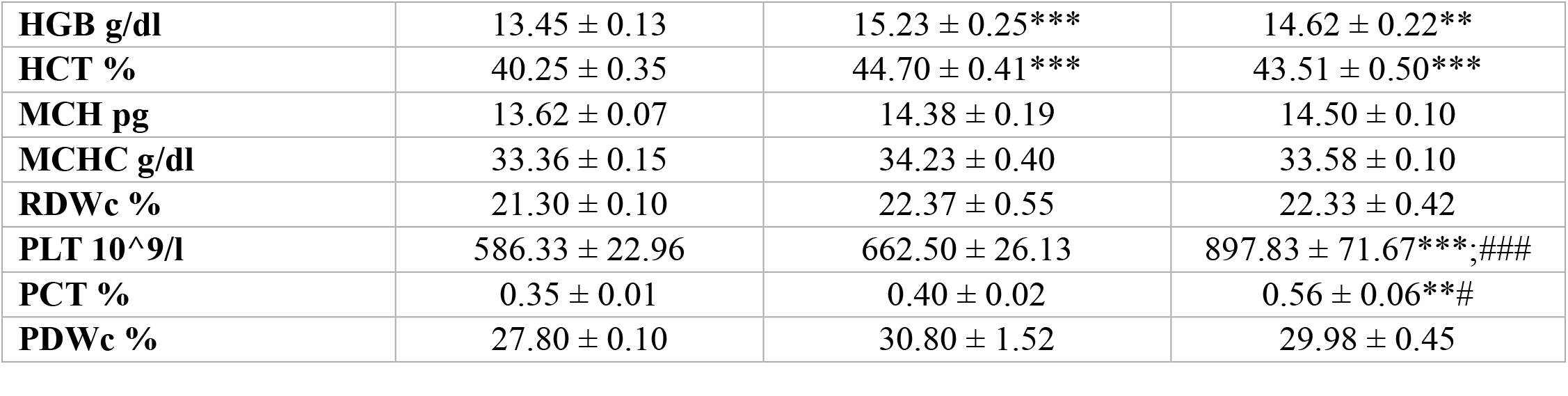
Serum biochemistry and whole blood parameters for WT, MiR^-/-^, and MDR^-/-^ mice. Data shown as mean ± SEM control. P values were calculated by one-way ANOVA with Tukey’s post-test. \**p*<0.05,\*\**p*<0.01 and \*\*\**p*<0.001. MiR^-/-^ versus MDR^−/−^ is denoted as # *p*<0.05 and ### p< 0.001.

### MDR deletion alters the transcriptional landscape of VAT

To uncover the molecular landscape underlying the metabolic alterations observed in MiR^-/-^ and MDR^-/-^ mice, transcriptomics was conducted using VAT from WT, MiR^-/-^, and MDR^-/-^ mice. Principal component analysis (PCA) showed that MDR^-/-^ mice were distinctly separated from WT mice, highlighting a significant genotype-specific transcriptional response. However, the PCA analysis showed that MiR^-/-^ mice overlapped and were located in proximity with WT mice, indicating only modest differences in their transcriptional profile (Figure S3). Volcano plots with a threshold of log_2_ fold change≥2 (log_2_FC≥2, FDR <0.05) identified differentially expressed genes (DEGs) comparing MiR^-/-^ and MDR^-/-^ mice to WT mice (Figure 3A).

Transcriptome analysis showed that MiR^-/-^ mice had far fewer changes than MDR^-/-^ mice, with only 9 upregulated genes and 5 downregulated genes (log_2_FC≥2 or ≤-2, FDR <0.05), suggesting a limited role of MiR in VAT (Figure 3). In contrast, MDR^-/-^ mice had a substantial transcriptional shift in their VAT, with 259 upregulated and 130 downregulated genes, highlighting profound VAT remodeling. For example, cholecystokinin (*Cck*) emerged as the most upregulated gene in MDR^-/-^ mice, with log_2_FC=12.6, FDR=7.19e-21 (Figure 3B). In obesity, *Cck* is upregulated by islet cells to compensate for obesity-induced insulin resistance (Lavine et al., 2010). Mice with *Cck* knockout are resistant to high-fat diet-induced obesity, suggesting that *Cck* is involved in metabolic regulation (Lavine *et al*., 2010); however, the role of *Cck* in VAT has not been defined so far. Interestingly, the elevated levels of *Cck* and leptin (*lep*) (both satiety signals) in MDR^-/-^ mice reflected a compensatory satiety response to positive energy balance, which may lead to weight gain in MDR^-/-^ mice. Additionally, *lep* may regulate the fatty acid synthase (*FASN*) expression in adipose tissue, since experimentally increased plasma *lep* concentrations in rats resulted in a decrease of *FASN* mRNA levels in fat (Mayas et al., 2010). Interestingly, while *FASN* is crucial for energy homeostasis and its expression is often higher in obese individuals, MDR^-/-^ mice showed a significant downregulation, while MiR^-/-^ mice showed a downregulated trend. This is interesting as in some obese individuals, *FASN* can be epigenetically downregulated (Sievert et al., 2021). It is possible that under severe metabolic stress or a hyperglycemic state, the body may have limited capacity to synthesize beneficial, insulin-sensitizing fats (Sievert et al., 2021). One recent study also showed that miR-15a and miR-16-1 target the 3′-UTR of *FASN* mRNA, and therefore miR-15a/16-1 markedly reduced endogenous *FASN* in breast cancer cells (Wang et al., 2016). Taken together, the transcriptome analysis highlighted that the gene expression is strikingly perturbed in MDR^-/-^ mice compared to the MiR^-/-^ mice alone, supporting the idea that the obese-insulin resistant phenotype is primarily driven by the whole MDR cluster.

Furthermore, Venn diagram visualization confirmed the above scenario (Figure 3C, 3D). Interestingly, only 1 gene in MiR^-/-^ mice and 251 genes in MDR^-/-^ mice were uniquely upregulated compared to WT; whereas only 7 upregulated genes were overlapping in both groups. These genes are primarily associated with obesity or metabolic dysregulation. Overall, MDR^-/-^ underscores gene trajectory driving the molecular signature contributing to obese-insulin resistant phenotype. A heatmap illustrating DEGs confirmed severely dysregulated metabolic phenotypes in MDR^-/-^ mice when compared to WT (Figure 3E, 3F). Comprehensive pathway and process enrichment analysis using Metascape did not reveal dysregulated pathways due to significant DEGs in MiR^-/-^ mice, except for upregulation of the cell division pathway (data not shown). In contrast, MDR deletion significantly upregulated genes, which were associated with mitotic cell cycle process, DNA replication checkpoint, and inflammatory response (Figure 3G). Conversely, downregulated genes were associated with metal ion transport, tight junction, hippo signaling pathways, and insulin signaling pathways (Figure 3H), suggesting elevated inflammation, dysregulated ion transport, obesity-induced adipose tissue fibrosis, and insulin resistance in MDR^-/-^ mice compared to WT mice. Furthermore, protein-protein interaction enrichment analysis identified the densely connected network components associated with DEGs in MDR^-/-^ mice (Figure. 3I, 3J). The molecular complex detection (MCODE) networks identified for individual upregulated (Figure. 3I) and downregulated (Figure. 3J) genes have been gathered to show the functional module of biological processes involved in the modulation of metabolic homeostasis. Furthermore, gene ontology (GO) (Figure S4) and Ingenuity pathway analysis (IPA) analysis (Figure S5) showed affected cellular components, biological processes, and molecular functions. In addition, cytoscape analysis identified the list of markers involved in the respective pathways (Table S1). Overall, MDR^-/-^ mice showed a significantly altered transcriptional landscape of VAT, suggestive of its critical role in adipose tissue-mediated metabolic regulation.

In summary, these findings underscore for the first time a causal role of cancer-associated complex clusters in metabolic processes and extend the significance of the MDR locus beyond oncological contexts. By linking this miRNA-lncRNA deficiency to extensive VAT transcriptional remodeling with new molecular pathways, excessive fat mass, impaired insulin sensitivity, inflammation, and a novel energy balance scenario, these findings lay the foundation for exploring MDR as a potential therapeutic target in obesity and related metabolic disorders.

### Limitations of the study and future direction

Collectively, these new findings provide the proof-of-concept that the absence of MDR contributes to severe metabolic aberrations with a more pronounced effect than the loss of miR-15a/16-1 alone. Even though our data strongly suggest that VAT plays a key role in this metabolic aberration, MDR may also interact with other tissues, suggesting that it may be involved in a more complex tissue specific role. Also, the use of human cells and tissues from obese and healthy subjects would provide invaluable physiological context and scope of MDR actions. Future work evaluating the roles of MDR in vitro and in vivo, extensively characterizing the various types of WATs and other metabolic tissues in mice and human, and the generation of tissue-specific knock-out models will be essential to fully elucidate its mechanisms and therapeutic potential.

## Resource availability

Data: Bulk RNA-seq data (raw fastq files) have been deposited to NCBI and are publicly available, SRA: PRJNA1498673. Any additional information required is available from the lead contact upon request.

Code: No original code was used in this manuscript.

## Acknowledgments

This work was supported by the Morris L. Lichtenstein, Jr. Foundation (M1500307). We thank Dr. Riccardo Dalla-Favera, Columbia University, for gifting the knockout mice. We also thank Professor Farida Sohrabji and Dr. Shameena Bake, Naresh K. Vashisht College of Medicine, Texas A&M University, for acquiring the MRI picture. We thank Texas A&M Preclinical Phenotyping Core for performing serum and blood analysis, Texas A&M Institute of genome Sciences and Society for mRNA sequencing, and VMBS Core Histology Laboratory for histology. We thank Navadeep Somashekar for his help in using Python. We also thank Kathryn Alise Kunz, Laura Packer, Raina Hummel, Erin Frances Hembrador Chua, Jhanvi Karthik, and Samuel Salazar for their help during mouse sacrifice.

## Author contributions

MC conceptualized the study. The experimental setup was designed and run by NS and SV. NS and SV were involved in formal analysis and visualization. MC, NS, and SV were involved in data interpretation. The manuscript was written by MC and NS. The supervision, project administration, and funding were provided by MC. All authors approved the final version of the manuscript before the submission.

