## Supplemental Information for "Uncovering a New Role of Dleu2/miR-15a/16-1 Cluster in Insulin Resistance and Obesity"

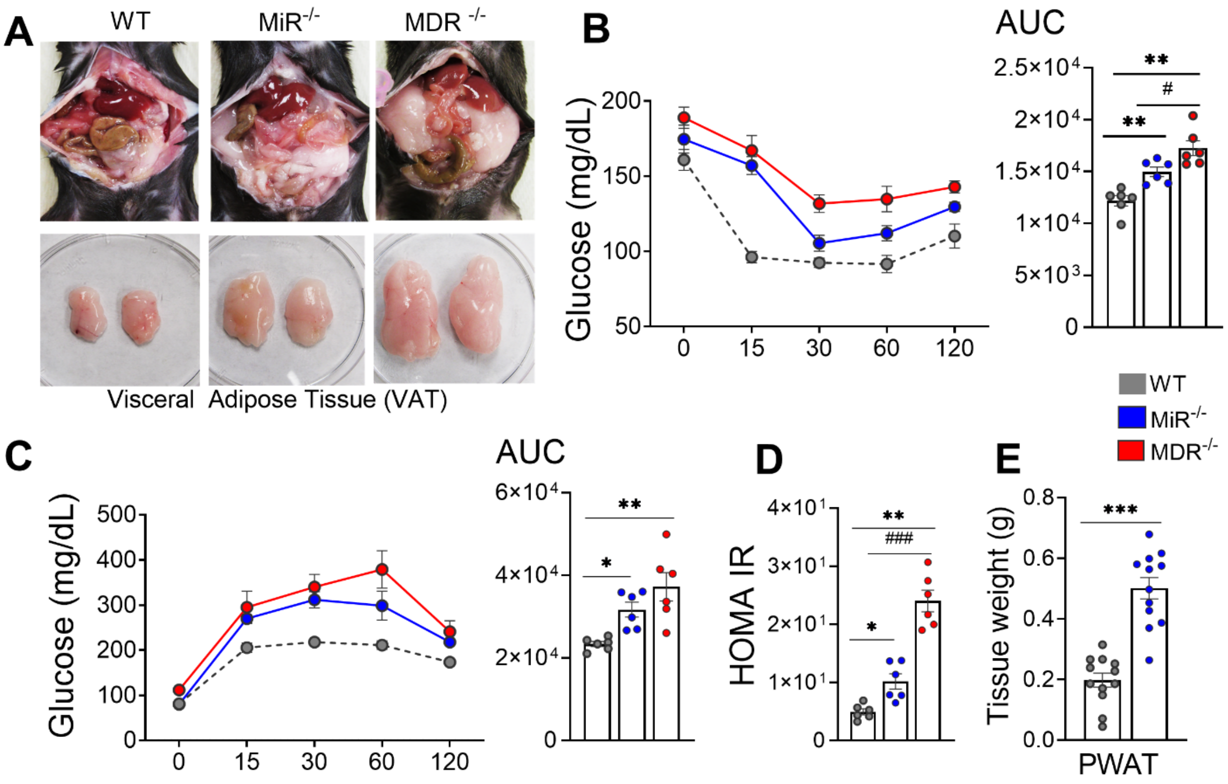


**Figure S1. MDR$^{-/-}$ contributes to excessive fat accumulation, leading to IR at 32 weeks of age.** (A) Representative image of the 32-week-old mice and corresponding VAT (*n*=12 per group). (B) Insulin tolerance test (*n*=6 per group) at 29 weeks, and quantification of the area under the curve (AUC). (C) Intraperitoneal glucose tolerance test (*n*=6 per group) at 30 weeks, and quantification of the area under the curve (AUC). (D) HOMA-IR.

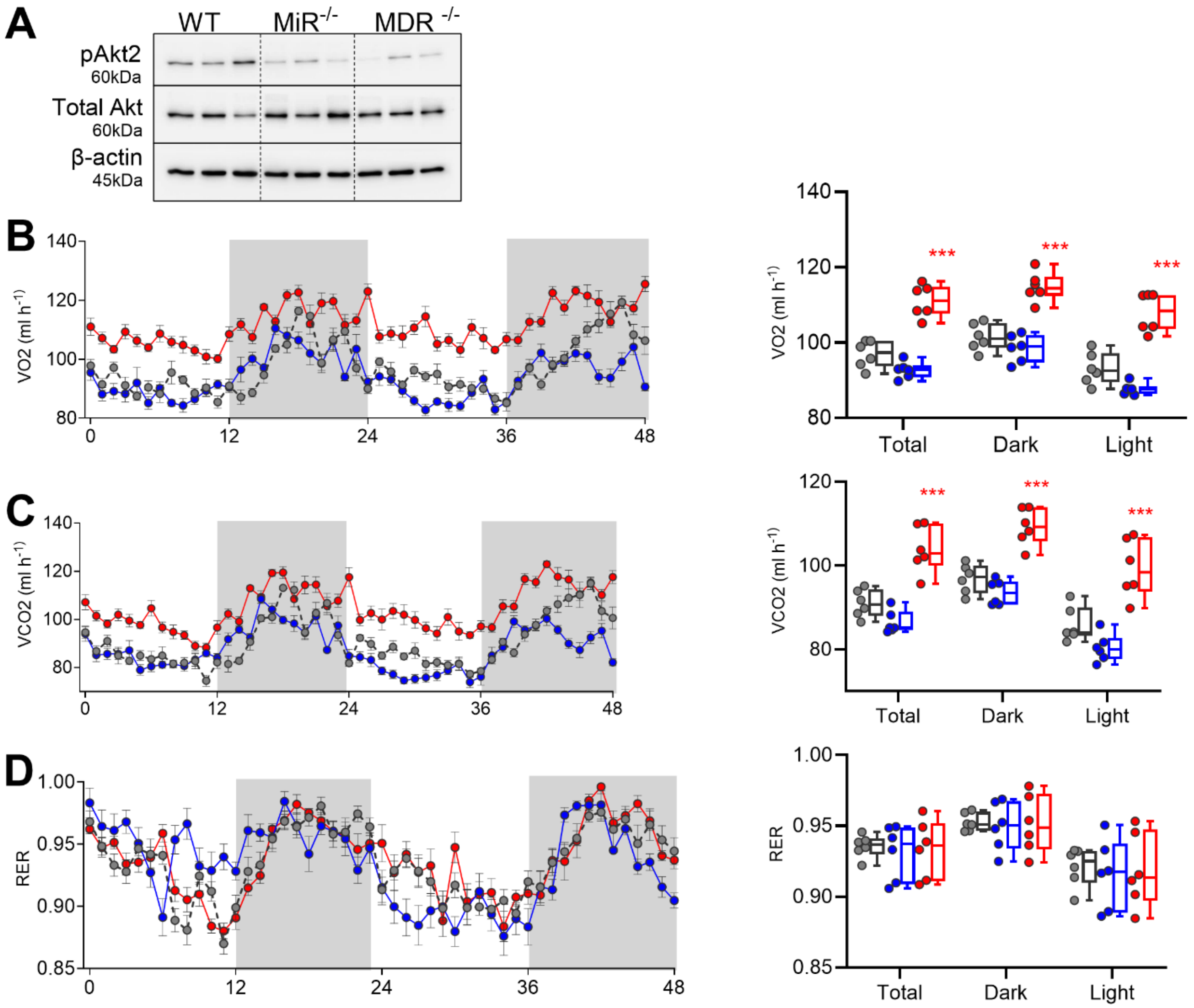


**Figure S2. MDR$^{-/-}$ contributes to higher $O_2$ consumption and $CO_2$ production.** Indirect calorimetry for 48h (*n*=6 per group), (A) Representative western blot image of insulin signaling marker (*n*=3 per group). (B) $VO_2$, (C) $VCO_2$, (D) RER. Line graphs (B-D) show representative 48 h trace means and error bars ± SEM, and bar graphs show 48 h means and error bars ± SEM.

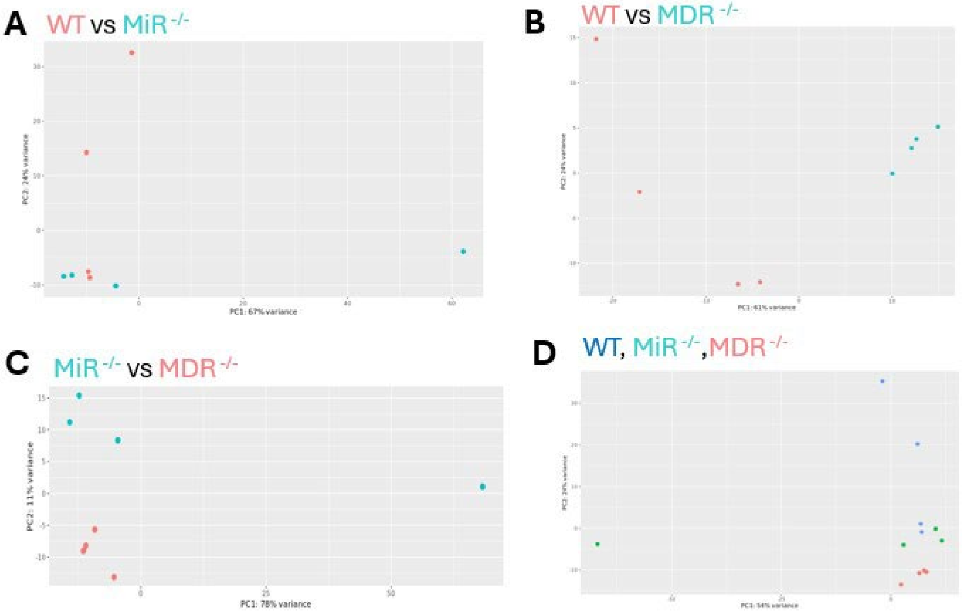


**Figure S3**. **PCA plots representing different groups clustering.** Each point represents one sample, and the color indicates groups: A) WT vs MiR $^{-/-,}$ B) WT vs MDR $^{-/-,}$ C) MiR $^{-/-}$ vs MDR $^{-/-}$ , D) WT mice, MiR $^{-/-}$ mice, and MDR $^{-/-}$ mice.

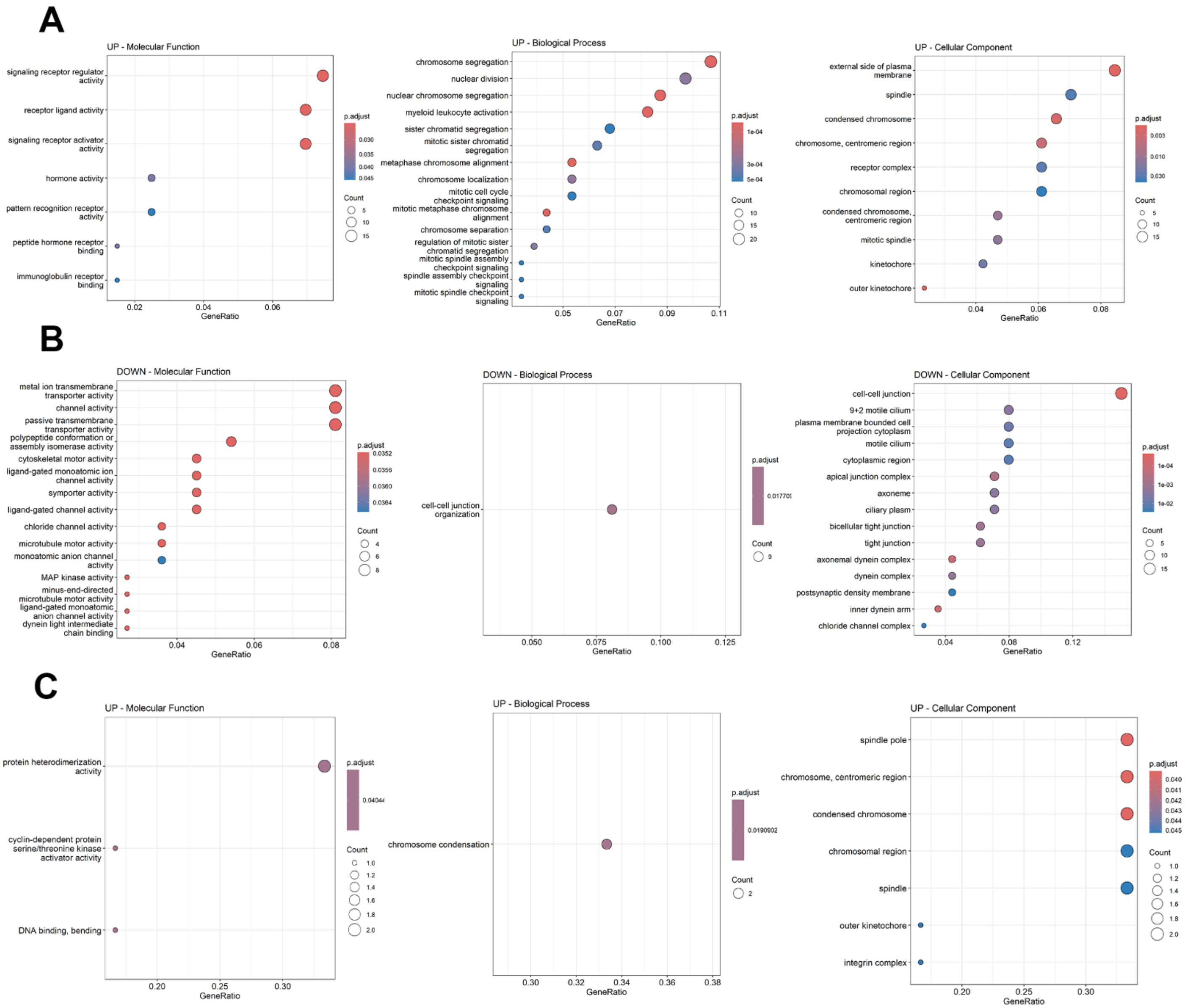


**Figure S4. Gene Ontology (GO) Analysis.** GO analysis identified the top molecular functions, biological processes, and cellular component. Circle size represents gene count and color indicates adjusted Padj value. (A) Downregulated genes in WT vs MDR$^{-/-}$, (B) Upregulated genes in WT vs MDR$^{-/-}$. (C) Upregulated genes in WT vs MiR$^{-/-}$.

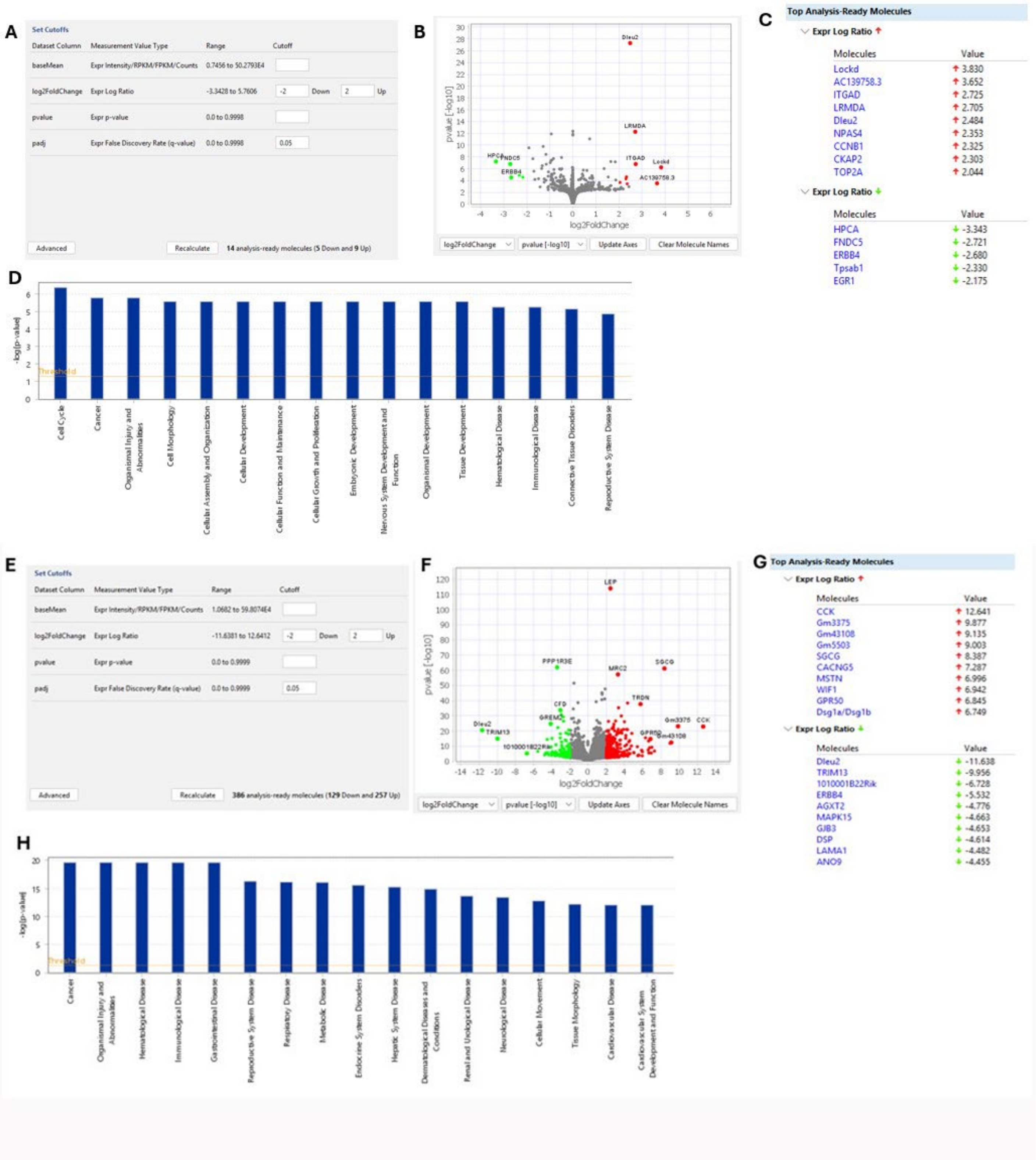


**Figure S5**. **Ingenuity Pathway Analysis (IPA)** of DEGs in WT vs MiR$^{-/-}$, (A) Set cut off. (B) Volcano plot. (C) Top DEGs. (D) Disease and Bio function. DEGs in WT vs MDR$^{-/-}$, (E) Set cut off. (F) Volcano plot. (G) Top DEGs. (H) Disease and Bio function.

### STAR Methods

#### Animals

Whole body miR-15a/16-1 and MDR knockout mice were a kind gift of Dr. Riccardo Dalla-Favera (Columbia University, New York). Mice were bred, and age matched mice were grouped and housed at ~22 °C with ad libitum access to standard chow diet (# 8604, Teklad™ Diet) and water throughout, following a 12:12 h light and dark cycle. Body weight was measured every week until 32 weeks of age. All procedures were performed in compliance with relevant laws and institutional guidelines and have been approved by the Institutional Animal Care and Use Committee of the Texas A&M Health Science Center (IACUC 2024-0191).

#### Metabolic Assessment

To assess glucose metabolism, mice underwent a glucose tolerance test (GTT). Briefly, overnight fasting mice were given a bolus intraperitoneal injection of Dextrose (# D9434, Millipore Sigma) in saline at 1.5 mg/kg body weight. Glucose level was measured in the blood at 0, 15, 30, 60, and 120 min using an Accu-Check, Contour®next glucose monitor. To evaluate the whole-body insulin sensitivity, an insulin tolerance test (ITT) was performed. After a 5 h fast, animals were injected with a bolus of Novolin® R (NDC 0169-1833-02, Novo Nordisk®) in saline to deliver 0.75 U/kg body weight. Glucose level during ITT was measured at the same time intervals as in GTT.

#### Noldus Open Field Test

The Noldus open field test was performed to examine spontaneous exploration behavior as described earlier. (Dominguez et al., 2005; Lai et al., 2006). One mouse at a time was placed in the center of an open field chamber of 40 cm × 40 cm dimensions, made of opaque acrylic (MazeEngineers, IL), and recorded under an overhead camera connected to the tracking software Ethovision XT (Noldus Information Technology Inc., VA) for 30 minutes. An automated tracking system then analyzed the footage for total distance traveled and time spent in the center. The center region was defined as the 10 cm × 10 cm square in the center of the chamber.

#### Body Composition

Mice were weighed, and whole-body composition was measured using the EchoMRI-100 whole-body composition analyzer (Echo Medical Systems, TX) to assess fat and lean mass as previously described (Wu et al., 2020).

#### Indirect Calorimetry

Metabolic activity in mice was measured using the PhenoMaster system (TSE-Systems, Germany) equipped to detect indirect calorimetry, measure food consumption, water intake, and monitor activity. Mice were placed into individual PhenoMaster cages and automatically monitored to record physiological parameters for 72h. During the first 24h of the acclimation period, the data collected was only used for validating the system setting, and the measurements over the next 48h were used for further data analysis. Room air went through the animal chambers at a rate of 0.5 L/min. Exhaust air from individual cages was sampled at 30-min intervals for 3 min. Sample air went through sensors to determine oxygen consumption ($VO_2$) and carbon dioxide production

($VCO_2$). Data expressed as the mean of 48h acquisition cycles. Data analysis was performed based on the recent guide to preclinical indirect calorimetry experiments (Banks et al., 2025) using Cal R software (version 2) and plotted in Graph Pad Prism (Version 11.0.1).

**Magnetic Resonance Imaging (MRI)**

The mice were imaged on a BioSpec 3-T MRI system (Bruker BioSpin, Germany). The whole-body MRI scans were recorded by acquisition with a standard T2 turbo spin-echo / turboRARE scan sequence. The fat tissue compartment was segmented into SAT and VAT. The VAT was defined as adipose tissue inside the abdominal cavity, excluding fat depots within abdominal and back muscles and fat tissue extending beyond the posterior outline of the vertebral body. The SAT was defined as subcutaneous fat external to the abdominal and back muscles.

**Dissections and Histology**

Animals were terminated by cervical dislocation. Blood was collected for serum biochemistry, and dissected tissues were weighed and snap-frozen in liquid nitrogen for RNA isolation or fixed in 10% paraformaldehyde (PFA) for 48h for histology. PFA-fixed adipose tissue pads were processed in 70% ethanol for at least 24h and embedded in paraffin. For hematoxylin and eosin staining, 4 µm thick sections of adipose tissue were stained by the standard protocol for measurement of the lipid droplet size. Images were taken at 10x magnification using a Nikon Microscope. Quantification was performed using Python (version 3.12.13), pyplot as plt from skimage import measure and segmentation feature. Three animals per group were analyzed, with 5 random area pictures per animal.

**Blood Chemistry**

Complete blood count was determined using DXC 700 AU Chemistry Analyzer (Beckman Coulter Ireland Inc., Ireland) at the Texas A&M Preclinical Phenotyping Core (TPPC). The test parameter and settings were followed by the test specification of each reagent.

**Serum Biology**

Blood was centrifuged at 4 °C at 10000 rpm for 10 min. Supernatant was collected and snap-frozen in liquid nitrogen. ELISA for Insulin (Crystal Chem Inc., # 62100) was carried out as per the manufacturer's instructions. ELISA kits for serum Adiponectin, Leptin, and TNFα (# MRP300, # MOB00B, # MTA00B, and # AF-401-NA R&D Systems, USA, respectively) were used according to the manufacturer's instructions.

**RNA-seq**

Total RNA libraries were constructed using the TruSeq Stranded mRNA Illumina kit. The RNA-seq experiment was performed by the Texas A&M Institute of Genome Sciences & Society (TIGSS) using the NextSeq 2000 sequencing systems (Illumina). Raw reads were adapter and quality-trimmed using Trim-Galore! (v0.6.10) with default parameters, including removal of Illumina adapter sequences and discarding bases with low Phred quality, and MultiQC (v1.30) was used to aggregate per-sample quality metrics and confirm adequate read quality. Pseudo alignment of reads to transcripts was carried out with Salmon (v1.5.1, RRID:SCR_017036). DESeq2 (v1.51.6) was used to identify differentially expressed genes (DEGs) between sample pairs using the R package (v4.3.2). Gene ontology analysis was performed using the Bioconductor package ClusterProfiler, org.Mm.eg.db, enrich plot, and ggplot. Metascape analysis was performed on DEGs to identify significantly enriched GO, KEGG, Reactome, and MSigDB terms and pathways, using hypergeometric testing with Benjamini-Hochberg correction. For the significantly upregulated and downregulated gene list, protein-protein interaction enrichment analysis was carried out using databases such as STRING, BioGrid, OmniPath, and InWeb_IM. The resultant network showed the subset of proteins that form physical interactions with at least one other member in the list. The Molecular Complex Detection (MCODE) algorithm was applied to identify densely connected network components. The MCODE networks identified for the individual gene list were gathered. The raw data obtained from the mRNA seq were normalized and uploaded to Ingenuity Pathway Analysis (IPA) (Qiagen Ltd.) DEGs were defined as $\log_2 FC \geq 2$ and $Padj<0.05$. Subsequent bioinformatics analysis was performed in IPA, including canonical pathway analysis, disease and function, regulator effects, upstream regulators, and molecular networks.

**Western Blotting**

Adipose tissues were homogenized using Qiagen Tissue Lyser in RIPA Buffer containing Protease and Phosphatase inhibitor; and ground tissue was suspended in NuPage LDS sample buffer (NP0008, Invitrogen) along with sample reducing agent (NP0009, Invitrogen) and heated for 5 min at 95 °C. Proteins were separated by SDS-PAGE and transferred to PVDF membranes. Those were then blocked with Licor Blocking Buffer for 1h at room temperature and probed with primary antibody overnight at 4 °C. After washing 3x with TBST, secondary antibodies were loaded for 1 h at room temperature. Primary antibodies used were Phospho-Akt (Ser473#9271, Cell Signaling Technologies) and β-actin (# 4970, Cell Signaling Technologies) at a 1:1000 dilution. Finally, membranes were developed by SuperSignal$^{TM}$ West Pico Plus chemiluminescent substrate (# 34580, Thermo Scientific, USA). Electrochemiluminescence (ECL) signals were imaged using Chemidoc (Bio-Rad Laboratories). Blots were stripped using Stripping Buffer (# 46430, Thermo Scientific, USA) and probed with mouse Akt2 (# 5239, L79B2, Cell Signaling Technologies, USA) at 1:1000 dilution. Secondary antibodies used were HRP Goat anti-rabbit IgG (# 926-80011, WesternSure, Li-Cor Biosciences, USA) and HRP Goat Anti-Mouse IgG (# 926-80010, WesternSure, Li-Cor Biosciences, USA) at a concentration of 1:5000. ECL signals were developed and measured as Phospho-Akt. Densitometry was performed using ImageJ software.

**Statistics and reproducibility**

Statistical analysis was performed using GraphPad Prism (version 11.0.1) and R (version 4.3.2). Statistical visualizations, including volcano plots, heatmaps, and Venn diagrams, were employed in Python (version 3.12.13). Key libraries included Pandas for data manipulation, Matplotlib and Seaborn for graphical rendering, and adjust text for label optimization.
